# NAPS: Integrating pose estimation and tag-based tracking

**DOI:** 10.1101/2022.12.07.518416

**Authors:** Scott W. Wolf, Dee M. Ruttenberg, Daniel Y. Knapp, Andrew E. Webb, Ian M. Traniello, Grace C. McKenzie-Smith, Sophie A. Leheny, Joshua W. Shaevitz, Sarah D. Kocher

**Author notes:** Correspondence: Scott W. Wolf < >, Sarah D. Kocher < >. These authors contributed equally (Dee M. Ruttenberg, Daniel Y. Knapp).

## Abstract

1. Significant advances in computational ethology have allowed the quantification of behavior in unprecedented detail. Tracking animals in social groups, however, remains challenging as most existing methods can either capture pose or robustly retain individual identity over time but not both.
2. To capture finely resolved behaviors while maintaining individual identity, we built NAPS (NAPS is ArUco Plus SLEAP), a hybrid tracking framework that combines state-of-the-art, deep learning-based methods for pose estimation (SLEAP) with unique markers for identity persistence (ArUco). We show that this framework allows the exploration of the social dynamics of the common eastern bumblebee (*Bombus impatiens*).
3. We provide a stand-alone Python package for implementing this framework along with detailed documentation to allow for easy utilization and expansion. We show that NAPS can scale to long timescale experiments at a high frame rate and that it enables the investigation of detailed behavioral variation within individuals in a group.
4. Expanding the toolkit for capturing the constituent behaviors of social groups is essential for understanding the structure and dynamics of social networks. NAPS provides a key tool for capturing these behaviors and can provide critical data for understanding how individual variation influences collective dynamics.

## Introduction

Massive advances in computing and imaging technologies enable new methods to study and quantify animal behavior. Recently, many pose estimation tools, such as SLEAP and DeepLabCut (Pereira et al., 2022; Lauer et al., 2022) have been developed to track the location of individual body parts of multiple animals as nodes on a skeleton (see Figure 1). With advancements in computational ethology, we are able to detect fine-grained behaviors, including grooming, touching, and gait changes (Berman et al., 2014, 2016; Gernat et al., 2018; Jones et al., 2020; Wiltschko et al., 2020; Geuther et al., 2021; Klibaite et al., 2022; Wang et al., 2022). Despite these advances, maintaining the identity of individuals in multianimal data (identity persistence) remains a challenge. This problem is especially pronounced when closely interacting animals may occlude each other’s body parts or when animals are partially or fully obscured by a complex environment. This problem is not always trivially solved with visual classifiers, as they struggle to classify morphologically similar individuals.

**Figure 1.**
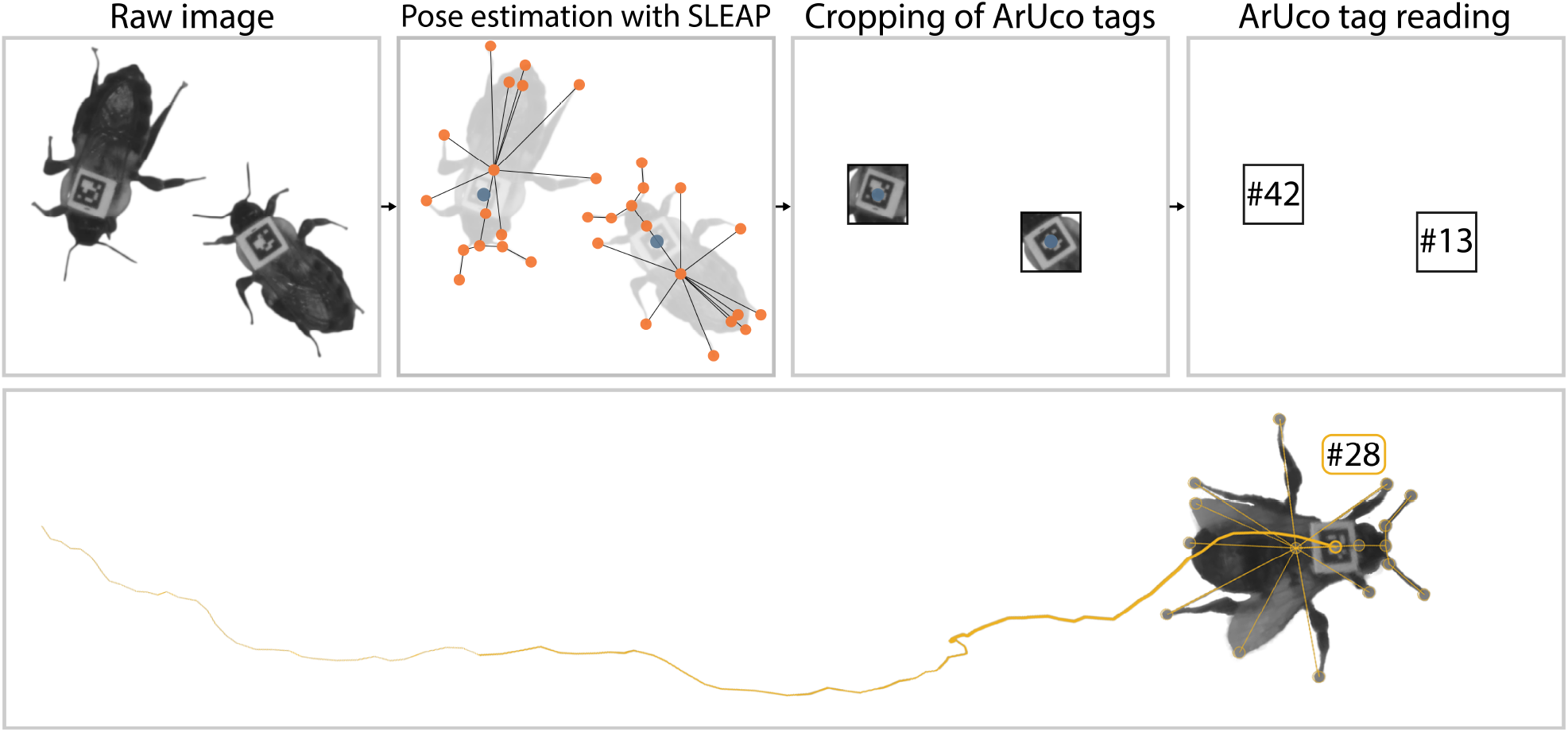
NAPS produces postural data with high-fidelity identity assignments from raw videos of tagged individuals. Top: Software pipeline. SLEAP is used to identify the location of each body part in the skeleton (orange dots) and the position of the ArUco tag (blue dot). NAPS crops the image around the tag and uses this as the input for ArUco tracking. Bottom: Tags are uniquely assigned to individual bees, and all instances from a single animal are combined to generate a unique identity. The orange line shows the tag’s location as the animal locomotes, increasing in width with time.

SLEAP, for example, provides two primary mechanisms for maintaining identity: temporal tracking, where identity is assigned based on the location of instances in the previous frame, and direct ID classification, which uses a neural network trained on known instances of a given individual to distinguish animals through differences in appearance. Temporal tracking is confounded by physical occlusion, and direct ID classification requires a significant time commitment for model training and struggles to distinguish visually similar individuals (Pereira et al., 2022; Lauer et al., 2022; Romero-Ferrero et al., 2019).

In parallel with deep learning methods for pose estimation, several unique identifier-based approaches (Mersch et al., 2013; Crall et al., 2015; Gernat et al., 2018; Boenisch et al., 2018; Alarcón-Nieto et al., 2018; Eagan et al., 2022) have been developed to maintain the identity of individuals. In these methods, a unique identifier affixed to an individual allows for tracking each individual’s position and orientation over time. While effectively addressing the identity persistence problem, many complex behaviors are not captured by these methods in isolation, as tag position only provides the position of a single point on the animal’s body.

Here, we introduce NAPS (NAPS is ArUco Plus SLEAP), a framework that combines the identity information from ArUco tags with temporal tracking and postural information from SLEAP (Figure 1). Combining deep learning-based pose estimation with unique markers for each individual enables long-term pose tracking of individuals with robust identity assignment. Here, we document this framework, provide an example use case, and discuss potential future applications.

### NAPS (NAPS is ArUco Plus SLEAP)

NAPS uses a hybrid tracking algorithm to provide high-dimensional behavioral data compatible with recently developed computational ethology techniques. Similar hybrid tracking techniques have been introduced in the literature (Smith et al., 2022; Gal et al., 2020). The NAPS framework utilizes quantitative information about body part position and unique identifying markers to provide postural data with identity over long timescales and robust to complex interactions. We used SLEAP, a deep learning-based framework for multi-animal pose estimation (Pereira et al., 2022), to localize the positions of body parts and the ArUco tags from video data. These body parts form the nodes of a user-defined skeleton (Figure 1). The output of the SLEAP workflow is a set of predicted body part locations combined into skeletons and their corresponding identities. However, SLEAP will split tracks from a single animal into multiple tracks if it does not correctly assign identity and can also propagate track-switch errors. We used the identity information from ArUco tags to link these broken SLEAP tracks together and remedy track switches (Figure 1).

The first step in our framework uses SLEAP, which can localize body parts with or without an identifiable tag. The SLEAP skeleton must include the tag as a node but otherwise can have as many other nodes as desired (Figure 1). To enable identification with ArUco, the area around the tag node is cropped out for each individual, and each crop is processed with OpenCV’s ArUco detection module (Bradski, 2000; Garrido-Jurado et al., 2014). We assign an identity to each instance using the ArUco tags through the Kuhn-Munkres algorithm (Kuhn, 1955, 1956; Munkres, 1957). The ArUco tag ID is read for each instance, generating a binary matrix of tag ID and SLEAP instance coincidences, *I*_*i,j*_(*t*) for frame *t*. To mitigate issues with misassigned and misread tags, we generate a cost matrix by subtracting these vectors in overlapping windows,

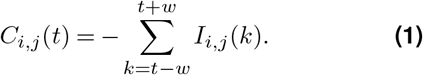

For the results shown here, we use a sliding window size of 41 frames (*w* = 20). We assign IDs to each instance by finding the minimum of the cost function using the Kuhn-Munkres algorithm as implemented in SciPy (Virtanen et al., 2020),

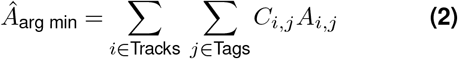

where *A* is a permutation matrix. Here, we only use the SLEAP-based tracking, i.e., forward propagation, for instances where the ArUco tag was read for the track previously and the ArUco-based ID was not reassigned to another track. This matching framework allows us to fix track switches and other errors produced by SLEAP when animals overlap in the image (Figure 2). The result is an easy-to-use system for combining the high-dimensional postural data from SLEAP with high-fidelity identity assignment from ArUco tags. This framework is embarrassingly parallelizable and thus scales easily to large datasets consisting of millions of images or more.

**Figure 2.**
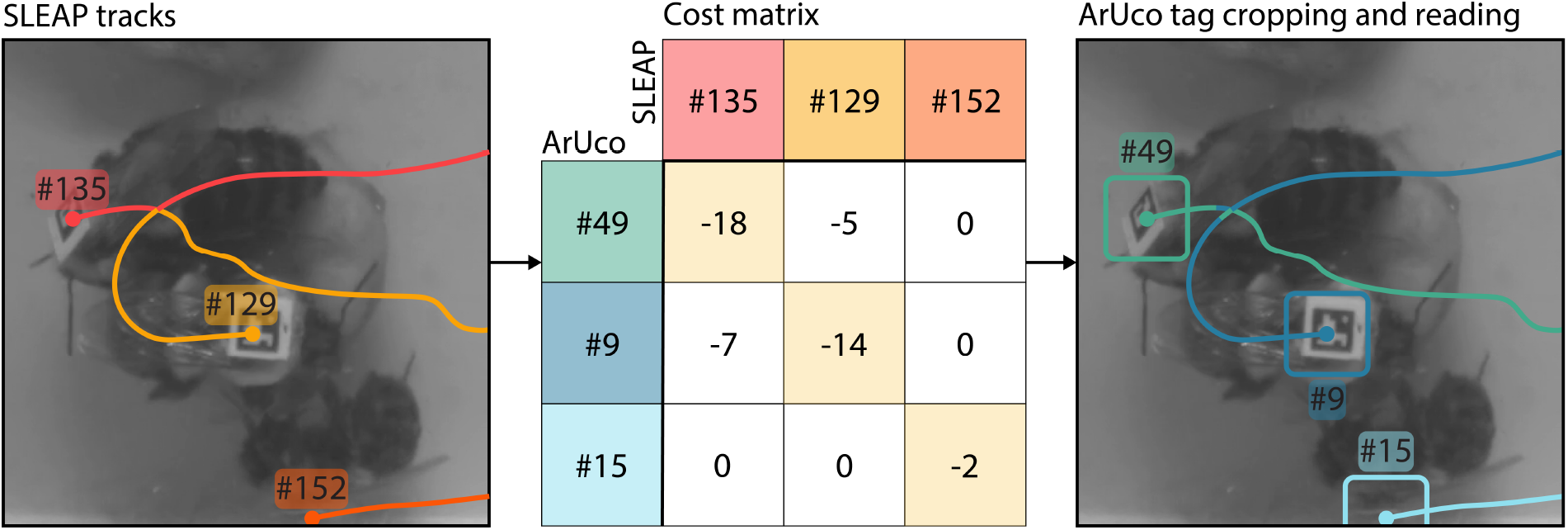
Left: Raw images of three interacting bees. Overlaid in color are the SLEAP-identified tracks that show a track-switch error. Middle: ArUco-based cost matrix, *C*, for this frame. Right: NAPS-corrected tracks overlaid without a track-switch error. The rounded boxes represent the area cropped out and subsequently read with OpenCV’s ArUco detection module for each instance.

### Example usage and assessment

As validation of our framework, we ran NAPS on an example dataset consisting of three one-hour video segments of 50 bumblebees (*Bombus impatiens*), including a single queen in a 235mm × 235mm arena (Figure S1 and Figure S2). Each hour segment was pulled from a 24-hour recording, with videos starting at 04:00, 12:00, and 20:00. The 50-bee colony was constructed from randomly sampled individuals, except the queen, from a colony obtained from Koppert Biological Systems (Howell, MI USA – August 2022) containing approximately 200 workers and a single queen. We printed 4.25mm × 5 5 ArUco tags generated from the 5×5_50 set on TerraSlate 5 Mil paper (TerraSlate, Englewood, CO USA) and cut these to 4.25 × 4.25 mm using a Silhouette Cameo cutting machine (Silhouette, Lindon, UT USA). Before tagging, bees were sedated via cooling on wet ice, and an ArUco tag was affixed to the dorsal side of the thorax using cyanoacrylate glue (Loctite, Hartford, CT USA) (Figure 1). As each tag was placed, we ensured that it could be read using a real-time ArUco tag reader and noted the tag number.

Following tagging, individuals were placed in arenas made from laser-cut acrylic and 3D-printed polylactic acid (PLA). Cotton wicks soaking in sugar water were placed in one corner of the arena to allow *ad libitum* feeding, and pollen mixed with honey was added for nutrition. The addition of these elements increases the complexity of the tracking environment and allows us to test NAPS’ performance under experimental conditions where its utility is most relevant.

Arenas were covered with clear acrylic and lit using two high-intensity 850nm LED light bars (Smart Vision Lights L300 Linear Light Bar, Norton Shores, MI USA) to allow continuous imaging of the bees while preserving a hive-like visual environment, as bees are unable to see infrared light (Figure S1). We imaged the arenas from above using a Basler acA5472-17um (Basler AG, Ahrensburg, Germany) camera recording 3664px × 3664px frames at 20 frames per second. Recordings were taken using campy to compress videos in real-time and drastically reduce file sizes (Severson, 2021). The videos have a spatial resolution of 15.5 pixels/mm, allowing us to capture fine-grained behaviors.

After video acquisition, we processed each video with SLEAP. We used a 17-node skeleton marking the head, two thorax points, abdomen, left and right antennal joints, left and right antennae, left and right wings, the pretarsus of each leg, and the ArUco tag. We trained on 31 frames sampled across the three videos. The 31 frames in our training set provided 1,550 labeled individuals, and the resulting model is highly accurate (see Figure S3). We performed inference using Nvidia A100 GPUs to generate initial tracks with temporal association. For this step, we use matching based on Intersection over Union (IoU) similarity with a 5-frame window. We also used SLEAP’s pre- and post-culling arguments, target instance count, and single track break connecting flag. The complete SLEAP command is provided in the Git repository (see Code and documentation). Following this, we ran NAPS on the dataset (for the entire command, see Code and documentation).

Finally, to compare the results from SLEAP, NAPS, and ArUco on our example dataset, we calculated both the percent of expected identifications (instances) per frame (Figure 3a) and the percent of realized identities (i.e., unique identities found) per video relative to the number of expected identities (Figure 3b). The number of expected instances identified in each frame varies by method, with SLEAP expected to identify all instances even when individuals are inverted or not active. ArUco can only capture instances in a given frame when the tag is identifiable, and the number of expected instances for this method is the number of bees which appeared active. NAPS will have the same number of expected instances per frame as ArUco but is able to capture instances detected by SLEAP but missed by ArUco in a given frame. We find that SLEAP is able to detect nearly all expected instances in each frame, NAPS captures almost as many instances as expected, and ArUco captures *∼*63% of the expected number of instances per frame (Figure 3a). The relative percent of expected identities assigned throughout a video also varies across methods, with SLEAP expected to detect all visible bees in a video, and NAPS and ArUco are only expected to detect bees that were active. SLEAP, using temporal tracking, assigns far more identities than expected, between *∼*156% and *∼*286% of the expected number (Figure 3b). NAPS and ArUco, with identities regulated by the detection of each ArUco tag, capture almost exactly the expected number of identities. These two metrics show that, while SLEAP is sufficient for identifying instances at the frame level and ArUco is sufficient for identifying the correct number of instances, NAPS enables the identification of the correct number of instances at the frame level while maintaining the correct number of identities across the video. Further, even within stretches of frames where equal numbers of identities exist, temporal identity tracking may cause identity swaps. In this case, NAPS will correct track switches to avoid the propagation of errors as shown in Figure 2.

**Figure 3.**
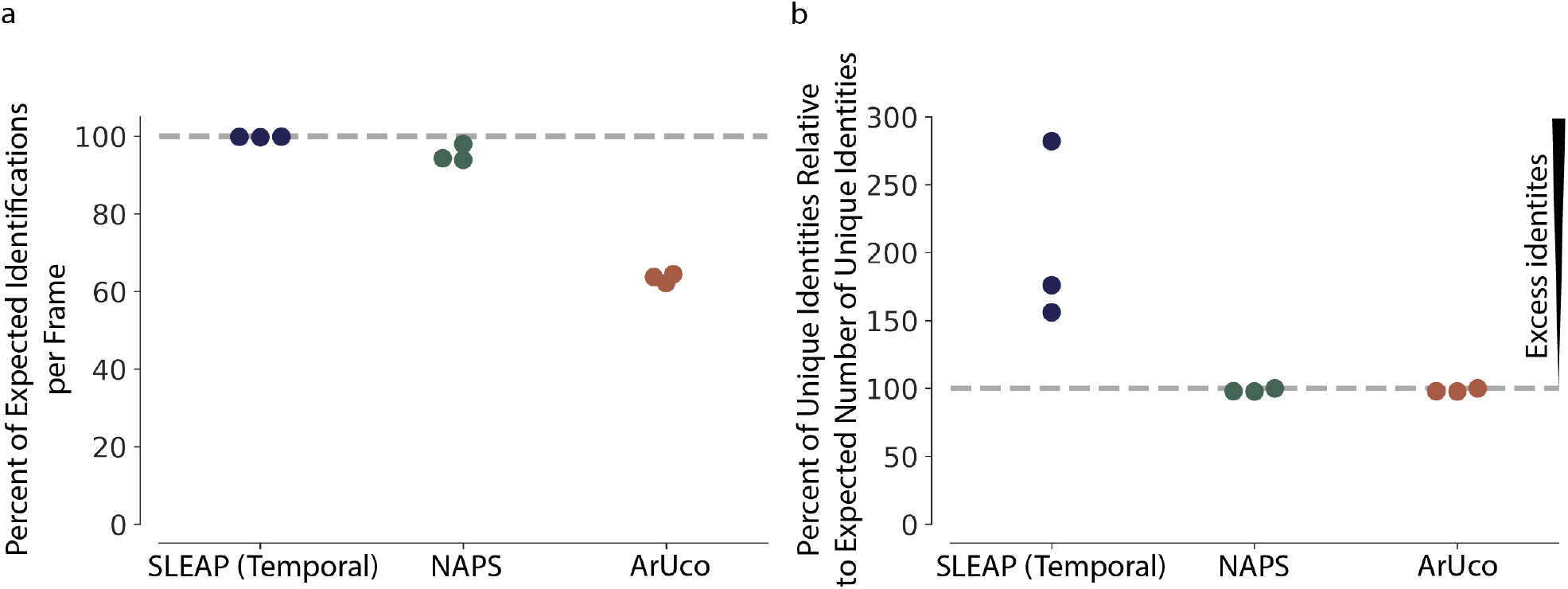
**a** A comparison of the percent of expected identifications per frame by tracking method (SLEAP, NAPS, and ArUco) in our example dataset. For our dataset, we expect all 50 bees to be identified by SLEAP. For NAPS and ArUco, we expect that all active individuals, where the ArUco tag should be detected at some point throughout the video, to be detected. This number varies from 44 to 48 depending on the video. Here, we see that NAPS captures a similar percentage of expected individuals per frame compared to SLEAP and significantly more than ArUco alone. **b** A comparison of the percent of the expected number of unique identities relative to the expected number of unique identities per video. We expect 50 unique identities for SLEAP and from 44 to 48 for NAPS and ArUco based on the number of active individuals in each video. SLEAP produces significantly more identities than expected while NAPS and ArUco generate almost exactly as many identities as expected.

The behavioral data from NAPS allows us to carry out detailed behavior analyses across dense groups (see Figure 4). For example, since bumblebees use their antennae for social interaction and communication, we labeled them as nodes in our SLEAP skeleton. This allows us to track antennal interactions across all pairs of bees in our example dataset (Figure 5a). Following the previous analysis from Wang et al. (2022), we quantified the amount of time each individual spent in different types of social interactions using a direct, convex-hull-based touch detector (Figure 5b). Bumble-bees can gain access to different chemical cues by antennating on different locations, and we can observe the time series of different modes of antennation (to the antennae, body, or abdomen) for a focal bee interacting with other bees in the arena (Figure 5c). By normalizing these interactions by their likelihood (Wang et al., 2022), we find that the antennae-antennae mode of touching is preferred in this large group of *∼*50 interacting bees (Figure 5d). This granular measure of an important behavior demonstrates how NAPS can extend the capacity of behavioral tracking platforms by quantifying previously difficult-to-measure behaviors of individuals in dense groups far larger than previously accessible.

**Figure 4.**
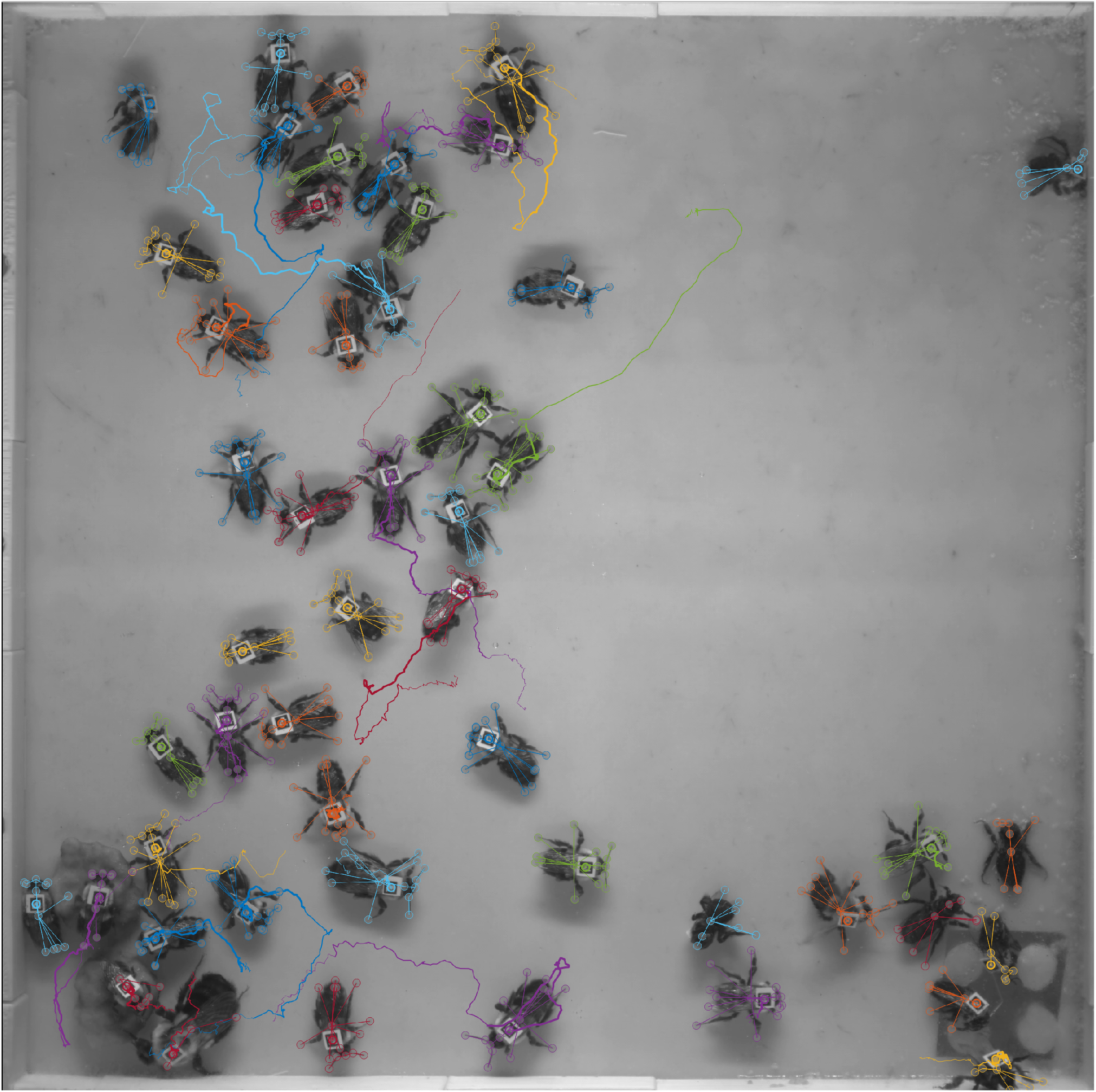
Example of a single frame from the output of NAPS. Traces for each individual, by color, show the animal’s location in the previous 250 frames (12.5 seconds). NAPS produces robust identity persistence and highly accurate localization of nodes for individuals. Here, all tracks are included in this image, including those not assigned to an ArUco tag.

**Figure 5.**
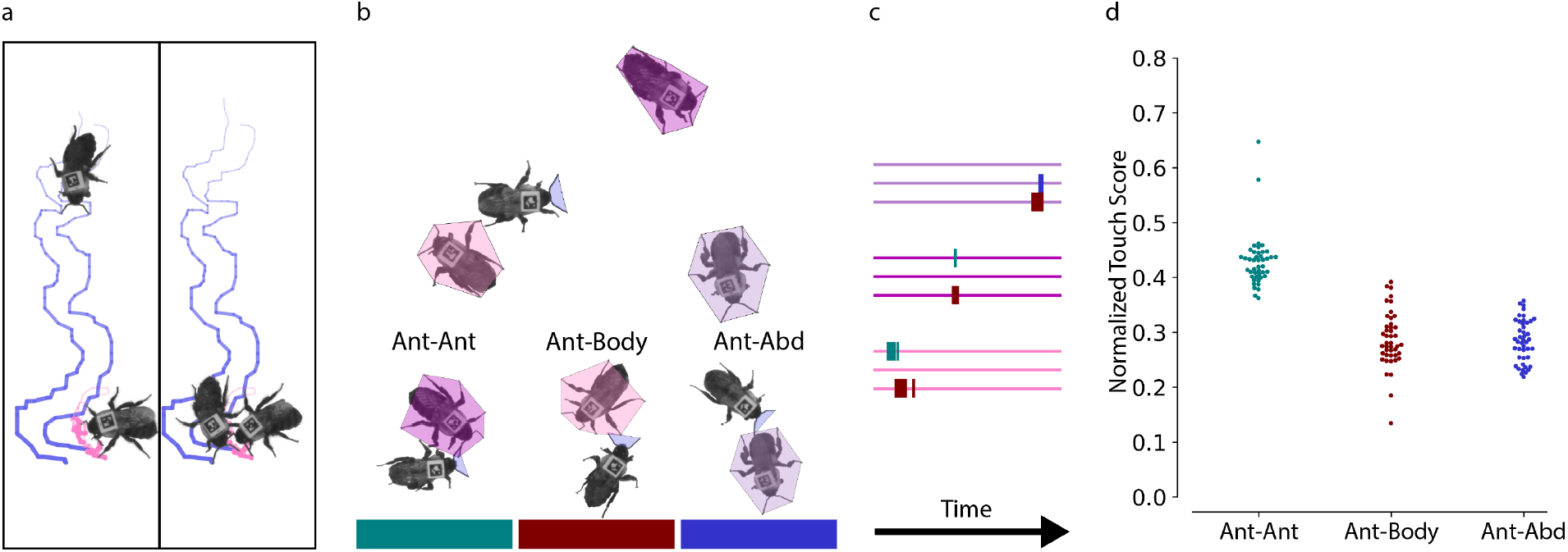
**a** Antennal tip traces of two bees during a brief interaction. The SLEAP coordinates of the antennal tips of two bees are plotted for 250 frames (12.5 seconds), providing spatiotemporal information about a single social interaction. **b** Schematic of antennal touch identifier (Wang et al., 2022). Instances of overlap between the antennal zone of the focal bee and the antennal (ant), body, or abdominal (abd) edges of the convex hull target bee are counted as touches. A bee can touch more than one of these zones at any moment. **c** Raster plot of antennal touches of the focal bee to three target bees over 400 frames (20 seconds). Touches to each target bee appear as color bars on the three raster traces associated with each target bee (color coded to match individuals in **b**), which each raster trace/color indicating a specific touch category. **d** Normalized touch scores across all bees with unique identities in all three sample videos (n=43 workers, one queen). For each touch category, the proportion of all touches for a given target bee, averaged across each video, is normalized to account for the different sizes of each body region and plotted by touch category (Wang et al., 2022).

## Discussion

Past studies of social behaviors utilized fine-grained behavioral data or persistent identity tracking with coarse behavioral tracking. NAPS combines these two types of data by integrating the pose estimation capabilities of SLEAP with the high-fidelity identity persistence of ArUco, enabling the concurrent analysis of behavior at both the individual and collective levels. This provides new opportunities to examine how individual behaviors vary in social contexts and the role of individual variation in shaping group and collective dynamics. Within a group, even genetically identical individuals may differ in their behaviors (Landeau and Terborgh, 1986; Herbert-Read et al., 2013; Bierbach et al., 2017; Werkhoven et al., 2021). With NAPS, users can probe how individual variation affects the dynamics of collective decision-making (Cook et al., 2020), the establishment of leadership and dominance (Pettit et al., 2015; Zanette and Field, 2009), how individuals adjust their behaviors in response to their local environment (Conradt and Roper, 2005), and the structure of individual and collective dynamics and their evolution (Hernández et al., 2021).

### Code and documentation

NAPS is an open-source framework. The source code is available on GitHub (github.com/kocherlab/naps), and the dataset used in this manuscript is available on Princeton’s DataSpace (doi: 10.34770/6t6b-9545).

To facilitate the broader usage of NAPS, we provide documentation, hosted at naps.rtfd.io, along with tutorials and example Jupyter notebooks (Brandl, 2022). We have released NAPS on Anaconda and PyPI, tested NAPS to nearly complete coverage (>94%), and new features are under continuous development. Further, we illustrate the usage of NAPS through Google Co-laboratory notebooks showing that the resources for a complete workflow, including SLEAP, are easily accessible using only free cloud computing resources.

While the NAPS framework relies on ArUco tags for individual identification, it can be directly extended to use any other type of visually distinguishable marker, such as color tags. This enables many different applications, including experiments where direct tagging may not be plausible. In parallel, NAPS users can take advantage of the flexibility of SLEAP, including the ability of end users to specify the nodes to track and capture previously elusive behaviors.

## Acknowledgements

We thank Emmanuel D’Agostino and Diogo Melo for thoughtful discussions, Eli Wyman for support wrangling bees, and Alexis Kane for helping with the generation of initial datasets. We further thank the Kocher Lab broadly for support with beekeeping and imaging.

This work was supported in part by an NIH Director’s New Innovator Award to SDK (1DP2GM137424-01), the Packard Foundation, the Princeton Catalysis Initiative, and the Department of Physics Undergraduate Research Fund at Princeton University. JWS acknowledges funding from NIH R01 NS10489. This work was also supported by the National Science Foundation through the Center for the Physics of Biological Function (PHY-1734030). SWW and DMR are supported by the NSF Graduate Research Fellowship Program (DGE-2039656), IMT is supported by the Lewis-Sigler Scholars Program at Princeton University, and GCM-S is supported by the Paul F. Glenn Laboratories For Aging Research at Princeton University.

## Authors’ contributions

SWW, DMR, JWS, and SDK conceptualized the project; SWW, DYK, and AEW developed the software; SWW, DMR, IMT, SAL, and SDK generated test data and beta tested the software; SWW, GCM-S, and SAL performed analysis on the example dataset. SWW and DMR drafted the initial manuscript. SWW, DMR, DYK, AEW, IMT, GCM-S, SAL, JWS, and SDK edited the manuscript and provided critical feedback. All authors approved the publication.

## Supplementary Information

### Figures

**Figure S1.**
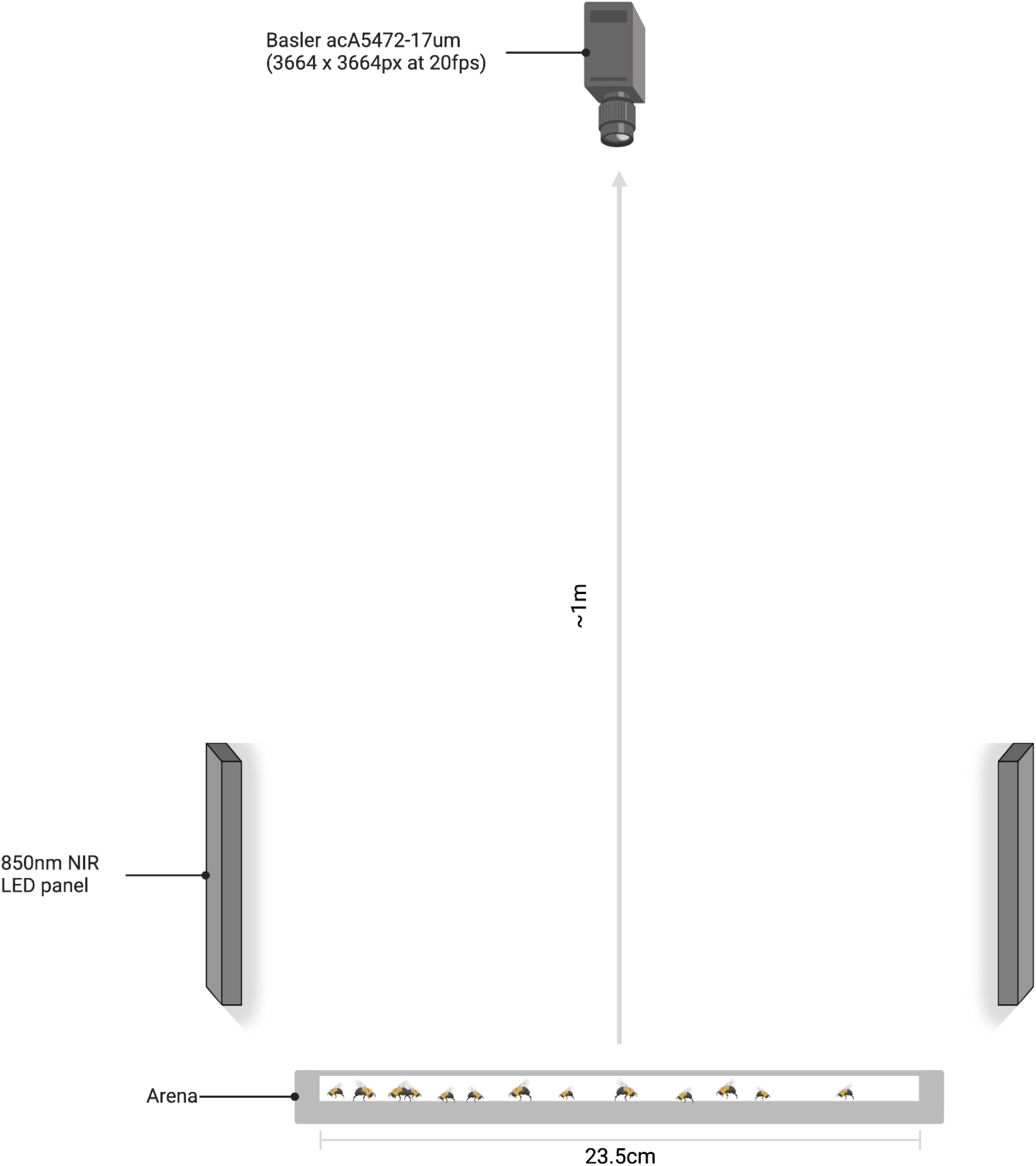
Diagram of the imaging setup used for capturing the example bumblebee data used. A camera facing directly down on the custom arena is flanked with two 850nm infrared LED light bars to allow continuous imaging of the bees without perturbation, as bees are unable to see infrared light. The resulting imaging setup allows high-quality imaging of large groups of individuals in a hive-like environment.

**Figure S2.**
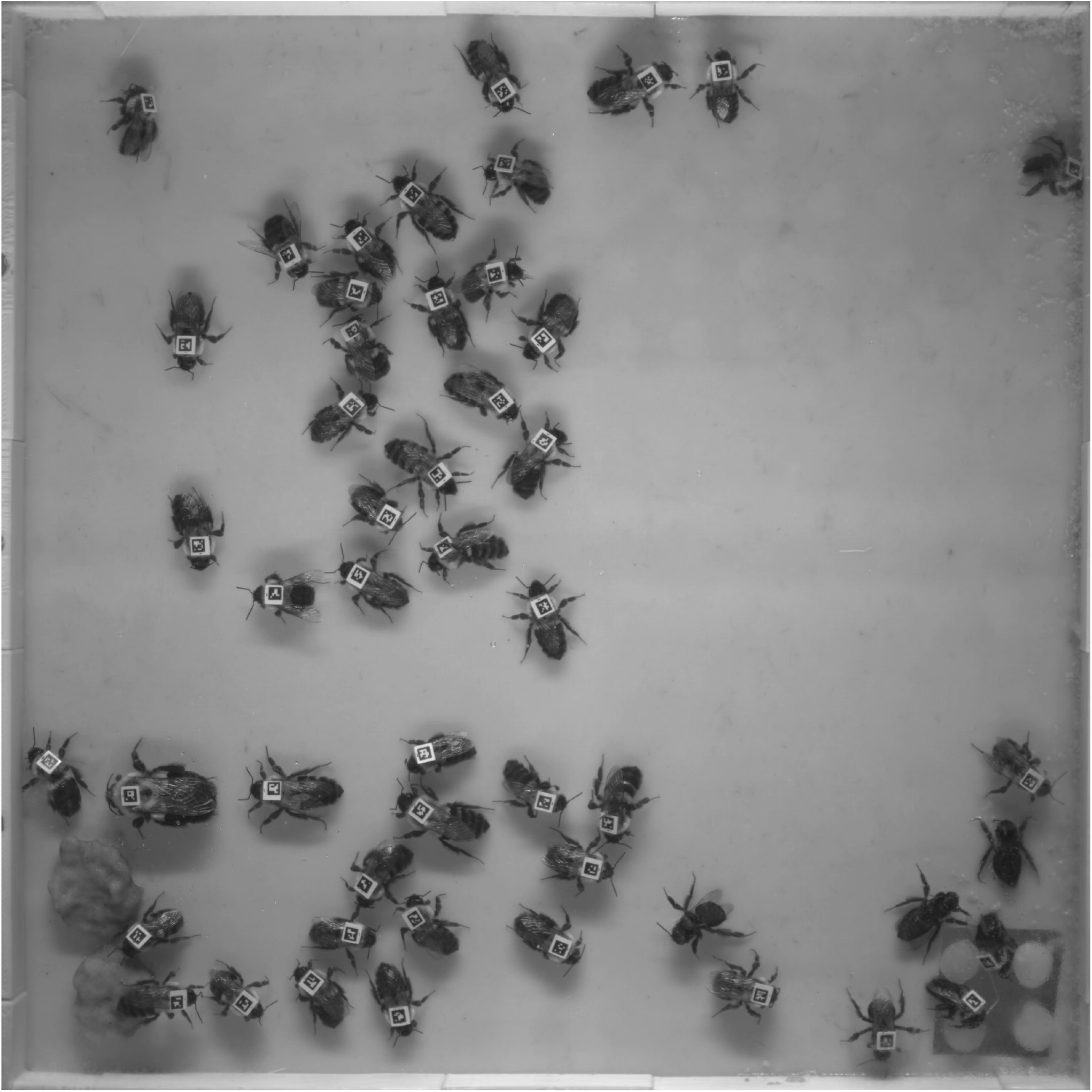
Example image pulled from the data set. Here, we see the density of individuals and the complex environment introduced by pollen dough (lower left) and sugar-water wicks for feeding (lower right).

**Figure S3.**
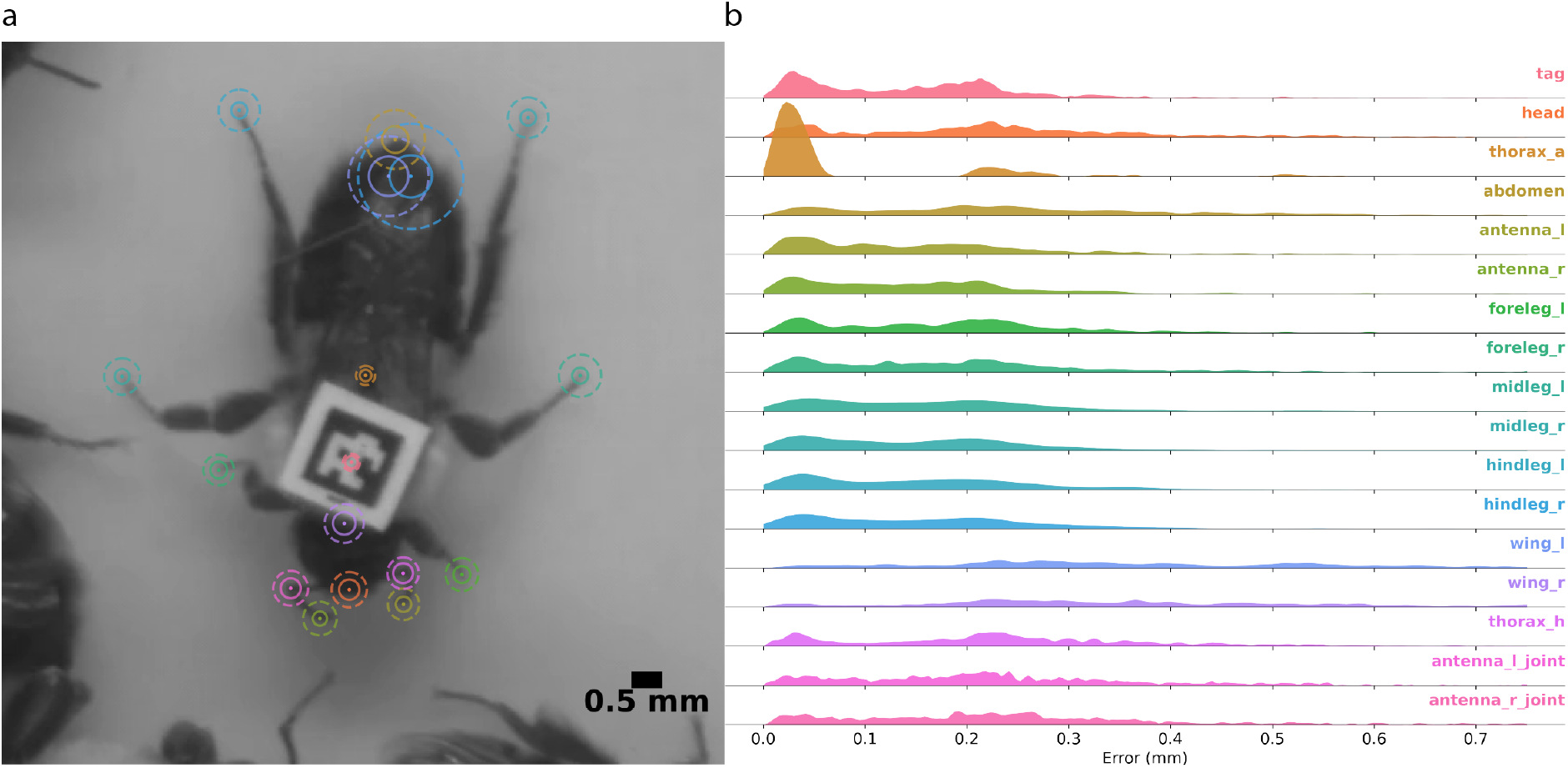
Node-wise localization accuracy. **a** shows the 75th percentile (solid) and 90th percentile (dashed) error for held-out validation set. **b** shows the distribution of localization errors on the same validation set. This distribution is clipped to (0, 0.75) for visibility.

